# Memory destabilization and reconsolidation dynamically regulate the PKMζ maintenance mechanism

**DOI:** 10.1101/2020.07.04.187823

**Authors:** Matteo Bernabo, Karim Nader

## Abstract

Useful memory must balance between stability and malleability. This puts effective memory storage at odds with plasticity processes like reconsolidation. What becomes of memory maintenance processes during synaptic plasticity is unknown. Here we examined the fate of the memory maintenance protein PKMζ during memory destabilization and reconsolidation. We found that NMDA receptor activation and proteasome activity induced a transient reduction in PKMζ protein following retrieval. During reconsolidation, new PKMζ was synthesized to restore the memory. Failure to synthesize new PKMζ during reconsolidation impaired memory but uninterrupted PKMζ translation was not necessary for maintenance itself. Finally, NMDA receptor activation was necessary to render memories vulnerable to the amnesic effect of PKMζ-antisense. These findings outline a transient collapse and renewal of the PKMζ memory maintenance mechanism during plasticity. We argue that dynamic changes in PKMζ protein levels can serve as an exemplary model of the molecular changes underlying memory destabilization and reconsolidation.

## Introduction

Long-term synaptic potentiation is stabilized by active molecular mechanisms that appear to maintain long-term memory in animals (Migues et al., 2010; Osten, Valsamis, Harris, & Sacktor, 1996; Sacktor et al., 1993). Considerable evidence shows that long-term memories can also become transiently labile following retrieval during a process known as memory destabilization. What triggers the switch from stability to plasticity is becoming increasingly clear with several initiating mechanisms identified (Ben Mamou, Gamache, & Nader, 2006; J. L. C. Lee & Flavell, 2014; Merlo et al., 2015; Suzuki, Mukawa, Tsukagoshi, Frankland, & Kida, 2008). Crucially though, how the pro-stability processes are affected and how they are recapitulated during reconsolidation remains unknown.

Following retrieval, memories can become temporarily unstable. This can be triggered by activation of a number of different receptors: N-methyl-D-aspartate receptors (NMDARs; Ben Mamou et al., 2006), cannabinoid receptors (J. L. C. Lee & Flavell, 2014), L-type voltage-gated calcium channels (Suzuki et al., 2008), or dopamine D1 receptors(Merlo et al., 2015). Destabilization also requires phosphorylation of CaMKII which promotes protein degradation (Jarome, Ferrara, Kwapis, & Helmstetter, 2016; A. S. Lee et al., 2008). Following a brief period of plasticity, the memory must be reconsolidated to ensure long-term stability. Importantly, infusion of protein synthesis inhibitors like anisomycin (Nader, Schafe, & Le Doux, 2000) or rapamycin (Jobim et al., 2012) after retrieval impairs long-term memory, suggesting that new protein synthesis is crucial to reconsolidation. Specifically which proteins must be degraded and synthesized remain largely unknown.

The length of the lability window is not clearly defined, with multiple biochemical processes occurring over different timescales. For instance, changes in α-amino-3-hydroxy-5-methyl-4-isoxazolepropionic acid receptor (AMPAR) content at amygdala synapses have been shown to occur as early as 5 minutes after retrieval and return to baseline by about 3 hours (Hong et al., 2013). Roughly 1 hour after reactivation of an auditory fear memory, there is an especially high concentration of phosphorylated GluA1-containing AMPARs at amygdala synapses compared to non-reactivated animals (Jarome et al., 2012). However by 6 hours, the lability window seems to be closed since protein synthesis inhibition can no longer disrupt long-term memory (Nader et al., 2000). Thus, destabilization of auditory fear memories appears to occur within 5 minutes of retrieval, persist for at least one hour, and reconsolidation seems to be complete by 6 hours after retrieval.

One critical component of this memory plasticity process is the transient reorganization of (AMPARs) at the synapse. During destabilization, GluA2-containing AMPARs are internalized and replaced with calcium permeable AMPARs (i.e. those lacking GluA2 subunits; Hong et al., 2013). During reconsolidation, the calcium permeable AMPARs are internalized and the GluA2-AMPARs are then reinserted into the membrane. While the movement of these receptors has been defined, what triggers their movement after retrieval is unknown. One clue lies in the regulation of GluA2-AMPA receptors by the protein PKMζ during memory maintenance (Migues et al., 2010).

PKMζ is an atypical protein kinase C (PKC) isoform, whose mRNA is transcribed from an internal promoter of the *Prkcz* gene (Hernandez et al., 2003). It contains the catalytic and hinge regions of the PKCζ protein but crucially lacks the regulatory domain of that protein (Sacktor et al., 1993). While there is still ongoing debate (A. M. Lee et al., 2013; Tsokas et al., 2016; Volk, Bachman, Johnson, Yu, & Huganir, 2013), research has primarily examined its potential role as a key regulator of long-term memory maintenance (Hardt, Migues, Hastings, Wong, & Nader, 2010; Hsieh et al., 2016; Sacktor et al., 1993; Shema et al., 2011). PKMζ seems to maintain long-term memories by regulating the movement of GluA2-containing AMPARs after LTP formation. It prevents endocytosis of GluA2-containing AMPARs (Migues et al., 2010) and limits their lateral diffusion (Yu et al., 2017). Given the role of PKMζ in GluA2-AMPAR trafficking, it is likely implicated in the AMPAR exchange that is central to memory destabilization/reconsolidation.

Here we investigated how memory maintenance, through PKMζ, is affected by destabilization and reconsolidation. We hypothesized that memory reactivation induces a transient disruption of maintenance mechanisms like PKMζ and that reconsolidation restores this mechanism (Fig. 1). We therefore tracked changes in PKMζ protein levels throughout this process and determined whether its synthesis is critical to reconsolidation. We found that destabilization induces an NMDAR-dependent reduction in PKMζ and that reconsolidation requires de novo protein synthesis to increase PKMζ and stabilize the memory.

**Figure 1.**
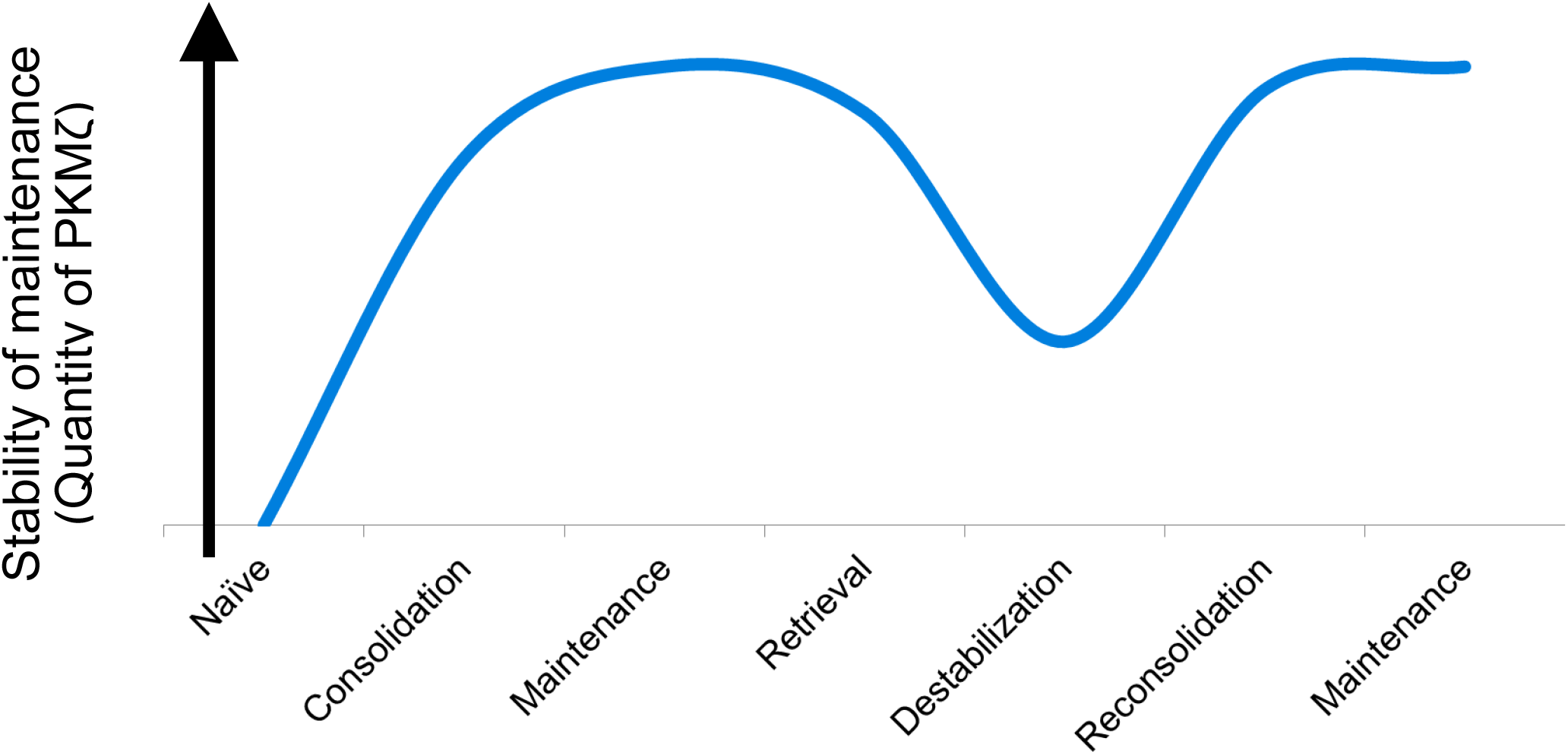
Hypothetical model of dynamic memory maintenance mechanism stability illustrated by changes in PKMζ expression. Consolidation of new learning increases expression of PKMζ from baseline levels. PKMζ expression remains relatively stable after consolidation as the memory is actively maintained. Retrieval triggers a reduction in PKMζ expression at the synapse while the memory undergoes destabilization. Reconsolidation increases expression of PKMζ, ensuring the long-term maintenance of the memory.

## RESULTS

### Experiment 1: Memory retrieval reduces synaptic PKMζ

While PKMζ has been much-studied in regards to memory maintenance, its response to memory retrieval remains unknown. Retrieval of conditioned auditory fear can induce internalization of GluA2-containing AMPARs in the amygdala (Hong et al., 2013). Therefore, we expected that a reduction in PKMζ in the BLA would be observed following recall of an auditory fear memory. Rats were trained by pairing a tone with a footshock. The next day, one group was exposed to one unpaired tone to trigger retrieval of the fear conditioning memory and sacrificed 1 hour post-reactivation. Rats in the second group did not undergo this re-exposure, remained in their home cage, and were sacrificed at the same time as the previous group. Western blots compared the PKMζ protein content in BLA samples of each group. Rats sacrificed one hour after the reactivation session showed less PKMζ than animals that remained in their home cage prior to sacrificing (Fig. 2A). Thus, memory retrieval seems to induce a decrease in synaptic PKMζ in the basolateral amygdala.

**Figure 2.**
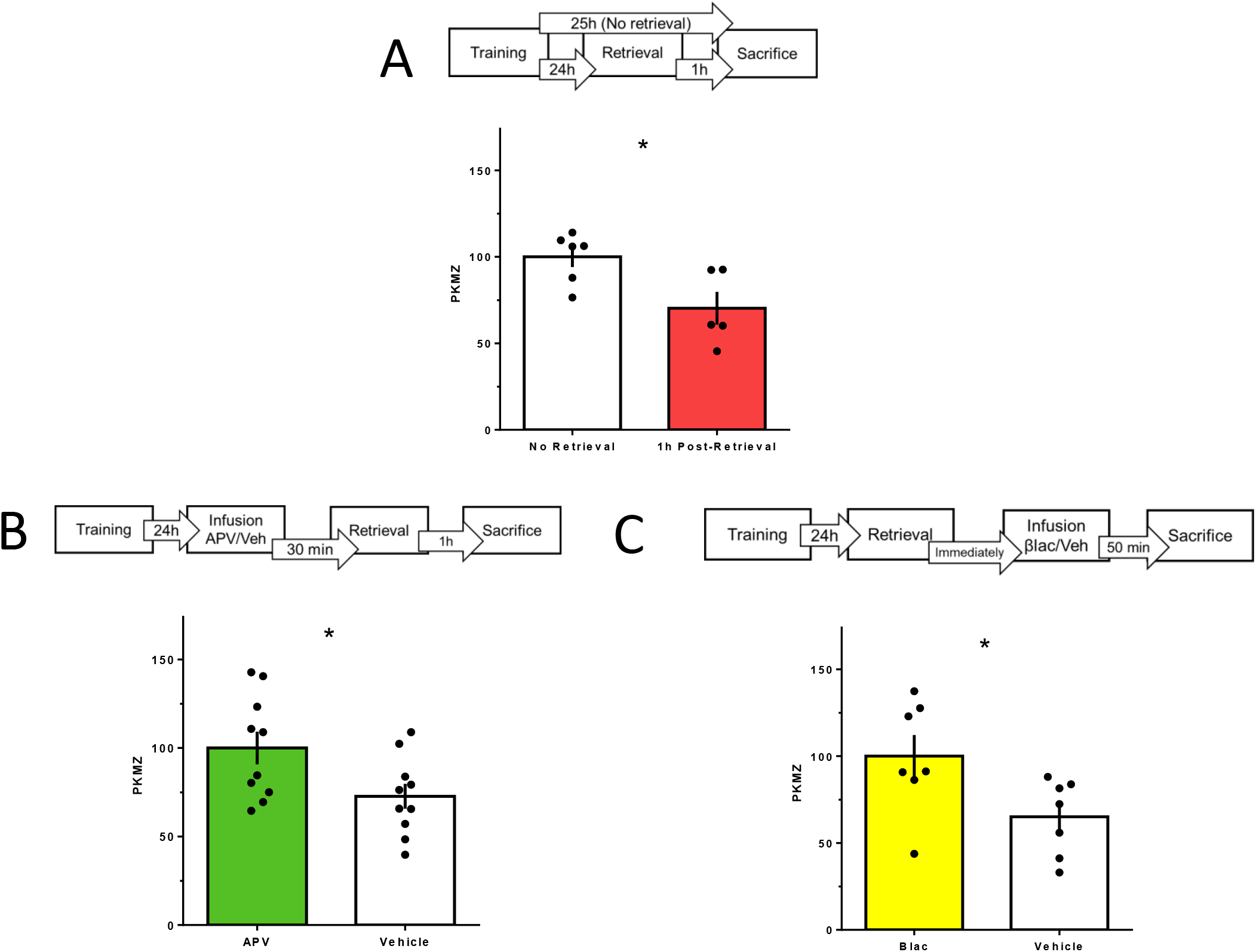
Downregulation of PKMζ during memory destabilization. **A.** Top: Following habituation, rats underwent auditory fear conditioning training in which a 30 second tone co-terminated with a 1.0 mA shock. Rats either experienced a retrieval test 24 hours later and sacrificed 1 hour post-retrieval or rats were sacrificed 25 hours after training (without retrieval). Bottom: PKMζ protein levels from Western blots. Rats sacrificed 1 hour post-retrieval showed significantly lower PKMζ protein in BLA tissue compared to rats that did not undergo retrieval (independent samples t-test, t_9=_2.755, p=0.022). **B.** Top: Rats underwent habituation and training. Thirty minutes prior to a retrieval test the next day, rats received an intracranial infusion of APV (5 μg/ μL in saline; 0.5 μL/side) or vehicle to the BLA. All rats were sacrificed 24 hours post-retrieval. Bottom: PKMζ protein levels from Western blots. Rats that received APV infusion prior to retrieval showed significantly higher PKMζ protein compared to rats that received vehicle (independent samples t-test, t_18=_2.358, p=0.030). **C.** Top: Rats underwent habituation, training, and a retrieval test. Immediately following retrieval, rats received an intracranial infusion of β-lac (32 ng/μL; 0.5 μL per side) or vehicle to the BLA. All rats were sacrificed 1 hour post-retrieval or roughly 50 min post-infusion. Bottom: PKMζ protein levels from Western blots. Rats that received β-lac infusion after retrieval showed significantly higher PKMζ protein compared to rats that received vehicle (independent samples t-test, t_12_=2.377, p=0.035).

### Experiment 2: Reduction in PKMζ is destabilization-dependent

Activation of NMDARs is necessary to initiate memory destabilization and internalization of GluA2-AMPARs (Ben Mamou et al., 2006; Hong et al., 2013). Therefore, we sought to determine whether the drop in PKMζ previously observed requires NMDAR activation. Rats were fear conditioned as before and, prior to retrieval, infused with APV, an NMDAR antagonist known to prevent labilization (Ben Mamou et al., 2006), or vehicle. Rats were then sacrificed one hour post-retrieval to compare PKMζ protein in the BLA. We found rats infused with APV did not show reduced levels of PKMζ seen in control animals (Fig. 2B). This suggests PKMζ protein decreases in response to NMDAR-dependent memory destabilization.

### Experiment 3: Proteasome activation is necessary for post-retrieval decrease of PKMζ

Following NMDAR activation, there is a CaMKII-dependent increase in proteasome activity (Jarome et al., 2016). Protein degradation seems necessary for destabilization since administration of the proteasome inhibitor β-lac prevents the amnesic effect of anisomycin on reconsolidation (A. S. Lee et al., 2008). Considering the outcomes of Experiments 1 & 2 showing that retrieval leads to a drop in expressed PKMζ, we therefore tested whether this decrease in PKMζ depends on proteasome activity. We infused β-lac or vehicle into the BLA of rats immediately following the retrieval of conditioned auditory fear. We sacrificed rats one hour after retrieval and compared synaptosomal PKMζ protein using Western blotting. We found that rats infused with vehicle showed less PKMζ after retrieval than rats infused with β-lac (Fig. 2C). Therefore, it seems that protein degradation following retrieval leads to the reduction in PKMζ we previously observed.

### Experiment 4: Reconsolidation increases synaptic PKMζ

Since destabilization reduces the availability of PKMζ at the synapse, we next tested whether reconsolidation involves a complementary increase in PKMζ protein. We re-exposed rats to one unpaired tone to retrieve the auditory fear conditioning memory and sacrificed them either 1 hour later (during destabilization) or 24 hours later (following reconsolidation). Western blotting showed that the quantity of PKMζ was higher in rats sacrificed 24 hours after retrieval compared to those sacrificed 1 hour after retrieval (Fig. 3A). This suggests that reconsolidation involves an increase of synaptic PKMζ to re-stabilize the memory.

**Figure 3.**
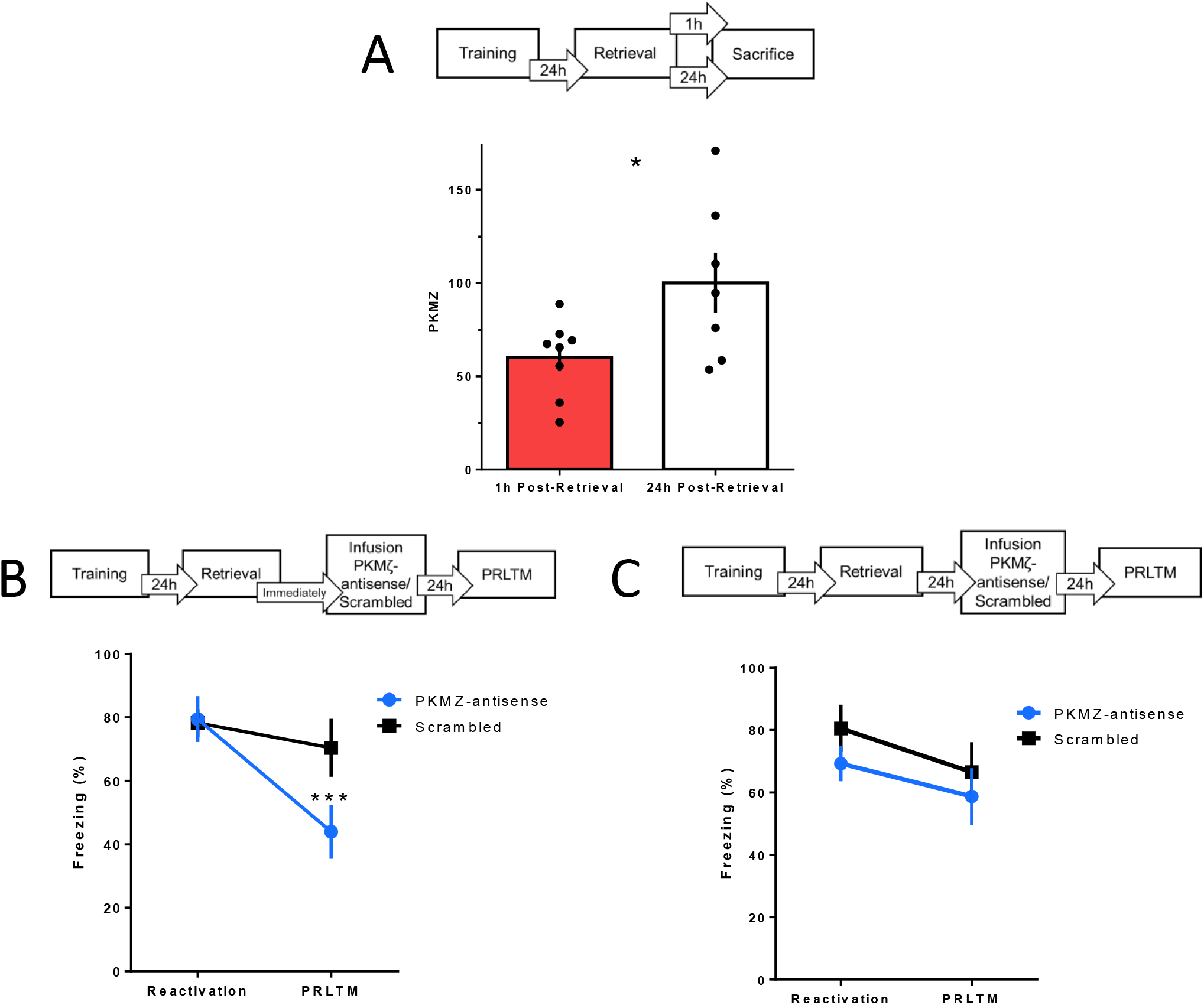
Increased expression of PKMζ via de novo synthesis during reconsolidation. **A.** Top: Rats underwent habituation, training, and a retrieval test and were sacrificed either 1 or 24 hours post-retrieval. Bottom: PKMζ protein levels from Western blots. Rats sacrificed 1 hour post-retrieval showed significantly lower PKMζ protein in BLA tissue compared to rats sacrificed 24 hours post-retrieval (independent samples t-test, t_13=_2.365, p=0.034). **B.** Top: Rats underwent habituation, and training. Immediately following retrieval, rats were infused with either PKMζ-antisense or scrambled ODN (2 nmol/μL; 0.5 μL/side). Rats were tested again 24 hours post-infusion. Bottom: Behaviour data showed a significant effect of the infusion on performance (one-way repeated measures ANOVA, F_1,18_=11.005, p=0.004). Rats that received PKMζ-antisense showed a significant impairment in performance when tested 24 hours post-infusion (Tukey’s test, p<0.001) whereas rats that received the scrambled sequence showed no impairment (Tukey’s test, p=0.210). **C.** Top: Rats underwent the same procedure as in B except that the infusion occurred 24 hours after the first retrieval test (rather than immediately afterwards). Bottom: Behaviour data showed no effect of infusion on performance (One-way repeated measures ANOVA, F_1,14_= 0.055, p=0.818).

### Experiment 5: Synthesis of PKMζ is necessary for reconsolidation

Given that reconsolidation increases the availability of synaptic PKMζ, we then investigated whether synthesis of new PKMζ is necessary to stabilize the memory. Immediately following a retrieval test, rats were infused with PKMζ-antisense oligodeoxynucleotides (ODNs) or a scrambled ODN sequence in the BLA. Rats were then tested 24 hours later to determine if PKMζ-antisense impaired reconsolidation and long-term memory retention. We found that rats’ memory differed depending on whether they were infused with PKMζ-antisense or the scrambled control (Fig. 3B). Specifically, rats receiving PKMζ-antisense showed impaired memory following infusion. However, rats receiving the scrambled control showed no difference in performance. Thus, these findings reveal that reconsolidation requires de novo synthesis of PKMζ.

### Experiment 6: Acute inhibition of PKMζ synthesis does not impair stable memory

Next, we evaluated whether transiently blocking PKMζ synthesis can induce a memory impairment on its own, or whether this effect requires memory retrieval. To this end, rats received BLA infusions with either PKMζ-antisense or a scrambled ODN 24 hours after a retrieval test. Thus, infusions occurred well after reconsolidation, which is believed to occur by 6 hours post-retrieval (Nader et al., 2000). Rats were tested again 24 hours after infusion. We found that rats’ freezing behaviour did not differ as a result of which ODN infusion they received (Fig. 3C). Neither group showed a memory impairment. Therefore, acute infusion of PKMζ-antisense after reconsolidation has already occurred does not disrupt long-term memory. Consequently, it seems that labile, but not stable, memories are vulnerable to disruptions of PKMζ synthesis.

### Experiment 7: Preventing destabilization protects memory from amnesic effect of PKMζ-antisense

Finally, we investigated whether it is specifically destabilization that renders memory vulnerable to PKMζ-antisense. That is, if we block NMDAR activation following retrieval, will PKMζ-antisense still impair retention of the retrieved memory? We infused rats prior to retrieval with either vehicle or APV to prevent memory destabilization. Immediately following retrieval, rats were then infused with either PKMζ-antisense or a scrambled control sequence and tested 24 hours later. We found that which drug (APV or vehicle) rats received prior to retrieval determined whether the post-retrieval ODN infusion could impair memory at the final test (Fig. 4). Rats infused with vehicle prior to retrieval followed by PKMζ-antisense showed impaired performance as described above. However, rats that received APV prior to retrieval followed by PKMζ-antisense did not show impaired memory. The scrambled control sequence did not affect memory in any group. This suggests that, absent destabilization, retrieved memories are not vulnerable to acute inhibition of PKMζ synthesis.

**Figure 4.**
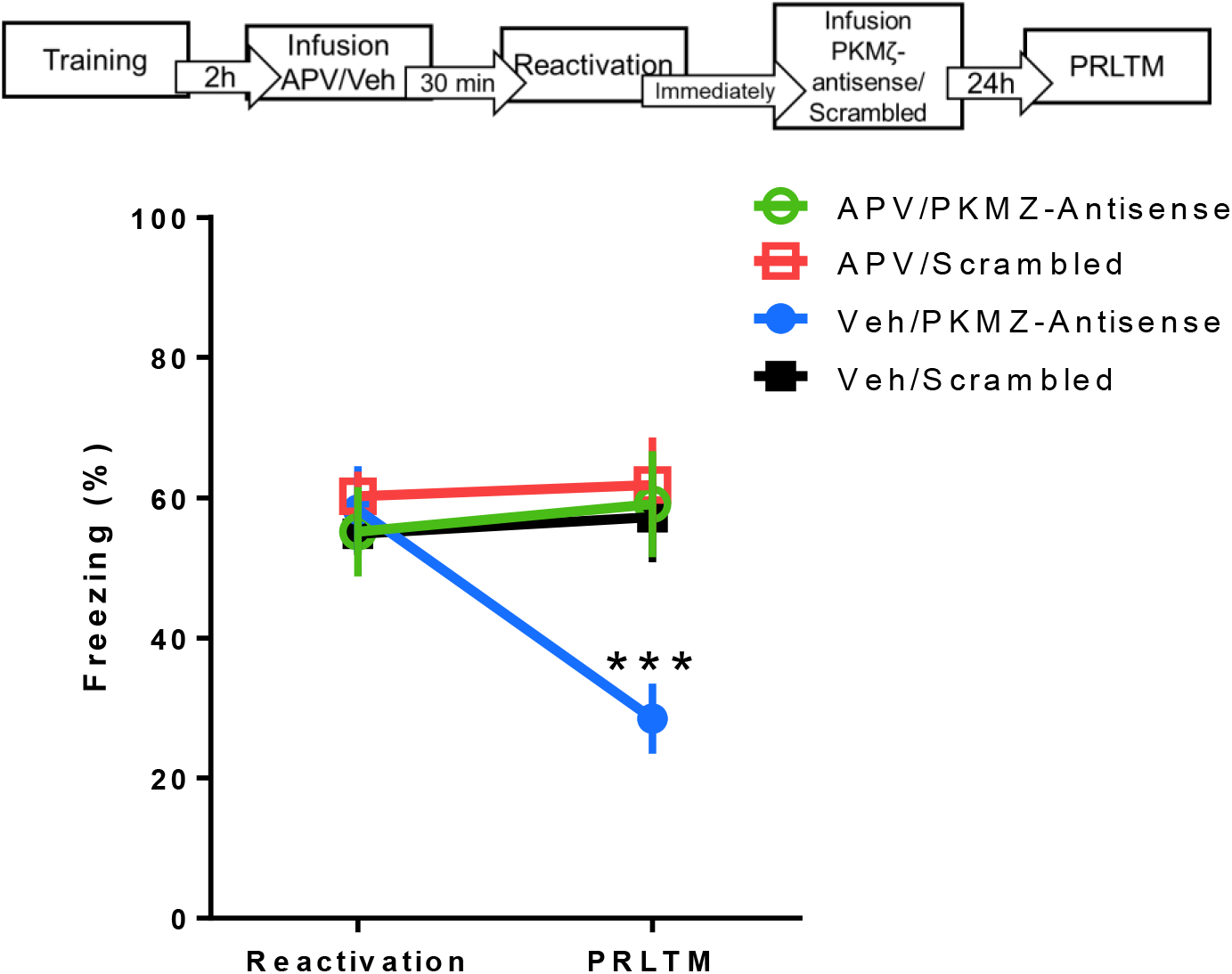
NMDAR activation renders memory vulnerable to impairment by PKMζ-antisense. Top: Rats underwent habituation and auditory fear conditioning training. One day after training, rats received an intracranial infusion of either APV or vehicle to the BLA, 30 min prior to a retrieval test. Immediately following retrieval, rats received an infusion of either PKMζ-antisense or scrambled ODN to the BLA. Rats were tested again 24 hours post-infusion. Bottom: Behaviour data revealed an interaction of pre-retrieval drug and post-retrieval infusion on performance (two-way repeated measures ANOVA, F_1,31_=6.65, p=0.015). That is, PKMζ-antisense had an amnesic effect in vehicle-treated animals (Tukey’s test, t_31_=5.045, p<0.001) but not in APV-treated rats (Tukey’s test, t_31_=−0.5428, p=0.999). Rats receiving the scrambled ODN sequence showed no impairment in performance.

## Discussion

Here we show dynamic regulation of PKMζ during memory destabilization and reconsolidation. After retrieval, there is a decrease in the availability of PKMζ at the synapse. This reduction requires both the activation of NMDARs and protein degradation. Within 24 hours after retrieval, PKMζ protein returns to the synapse. Indeed, de novo synthesis of PKMζ is necessary for reconsolidation and without it, long-term memory is impaired. Importantly, transient disruption of PKMζ synthesis outside of the lability window does not disrupt long-term memory. In fact, without NMDAR activation memories are not rendered labile after retrieval and acutely blocking translation of PKMζ does not impair memory.

To date, synthesis of at least four proteins, zif268 (Barnes, Kirtley, & Thomas, 2012; J. L. C. Lee, Everitt, & Thomas, 2004), Arc (Maddox & Schafe, 2011), C/EBPβ (Milekic, Pollonini, & Alberini, 2007), and C/EBPδ (Arguello et al., 2013), has been shown to be necessary for reconsolidation. That these proteins must necessarily be synthesized is perhaps not surprising. Each plays a role in gene expression which has repeatedly been shown to be crucial for reconsolidation (Nader et al., 2000; Tronson & Taylor, 2007; Villain, Florian, & Roullet, 2016). On the other hand, the predominant role of PKMζ seems to be memory maintenance. Therefore, this work provides evidence for a specific synaptic function that is disrupted and restored during destabilization and reconsolidation, respectively.

These findings are in line with previous research showing reorganization of AMPARs during memory destabilization and reconsolidation. During destabilization, GluA2-containing AMPARs are internalized (Hong et al., 2013) at the same time that we observed reductions in PKMζ expression. Further, the return of these AMPARs (Hong et al., 2013) coincides with an increase in PKMζ.

In recent years, the importance of PKMζ to memory maintenance has been debated since PKMζ-null mice show intact learning memory and are vulnerable to memory impairment by ζ-inhibitory peptide (ZIP). ZIP has traditionally been used to disrupt PKMζ catalytic activity (Kwapis, Jarome, Lonergan, & Helmstetter, 2009; Ling et al., 2002; Migues et al., 2010; Serrano et al., 2008) although it is also capable of inhibiting the activity of another atypical PKC isoform, PKCι/λ (Bogard & Tavalin, 2015; Ren et al., 2013). However, studies using more specific techniques have supported the role of PKMζ in memory. Overexpression of PKMζ enhances memory (Shema et al., 2011; Xue et al., 2015). On the other hand, expression of a dominant negative PKMζ one week after training impairs conditioned taste aversion memory (Shema et al., 2011). Administration of PKMζ-antisense has been shown to disrupt long-term memory in vivo (Hsieh et al., 2016) and late-LTP in vitro (Tsokas et al., 2016). Further, recent work showed that PKMζ-knockout mice may in fact compensate using another atypical PKC isoform, PKCι/λ, to maintain long-term memory in the absence of PKMζ (Tsokas et al., 2016). Nonetheless, to avoid the identified complications of ZIP, we limited our tools to antisense ODNs which offer sequence specificity and Western blotting. Thus, we feel confident that our results reflect the dynamics of PKMζ and not another kinase.

One limitation of this work is the uncertain fate of PKMζ post-reactivation. That PKMζ protein is reduced in the synaptosome could mean that it is being trafficked elsewhere or perhaps degraded. Our results do show that proteasome activation is necessary to reduce PKMζ post-retrieval. However, it is not clear whether PKMζ itself is degraded or if this effect is simply downstream of the degradation pathway. The requirement that new PKMζ be synthesized for reconsolidation could mean that existing PKMζ stores are actively degraded and must be restored, although this is speculation at present. Given the role of KIBRA in preventing degradation of PKMζ (Vogt-Eisele et al., 2014), it may be that reactivation triggers a decoupling of KIBRA and PKMζ leading to its degradation. Thus, future research should investigate the interaction of KIBRA and PKMζ during destabilization.

Numerous studies have found that acute inhibition of PKMζ kinase activity causes long-term impairment of memory (Gámiz & Gallo, 2011; Hardt et al., 2010; Shema, Sacktor, & Dudai, 2007). We found that transiently blocking the synthesis of PKMζ at 24 hours post-retrieval does not impair memory one day after infusion. This may reflect an oversaturation of PKMζ protein at the synapse, beyond what is required to maintain the memory. It may also indicate a long half-life for PKMζ, allowing existing protein to overcome a temporary reduction in new translation.

Altogether, this work shows that memory destabilization and reconsolidation involve the disruption and recapitulation of a crucial memory maintenance process. That is, plasticity occurs at the expense of PKMζ and restoring stability requires the production of new PKMζ. In the future, other critical proteins, both to induce lability and to restabilize synapses, will need to be identified in order to gain a full understanding of memory plasticity.

## Methods

### Animals

Male Sprague-Dawley rats (275-300g) were obtained from Charles River, Saint-Constant, Quebec. Rats were housed in pairs and maintained on a 12h light/dark cycle (lights on at 7:00 am, lights off at 7:00 pm). Rats received food and water ad libitum. All methods and procedures were approved by McGill University’s Animal Care Committee and conformed to Canadian Council on Animal Care’s guidelines.

### Surgery

Rats were given an intraperitoneal injection of anesthetic cocktail (1 mL/kg) containing ketamine (50 mg/mL), xylazine (3 mg/mL), and dexdomitor (0.175 mg/mL). Prior to surgery rats also received carprofen analgesic (5 mg/mL; 1 mL/kg) subcutaneously. Rats were bilaterally implanted with 22-gauge guide cannulas targeted to the basolateral amygdala (from bregma: AP −3.0 mm; ML +5.3 mm; DV −8.0 mm). Cannulas were secured to the skull with dental cement and three jeweller’s screws. To ensure the interior of the cannula remained clear of debris, metal dummies were inserted and remained in place except during infusions. Following surgery, rats were given an intraperitoneal injection of anesthetic reversal containing 0.5 mg/mL of antisedan. Following surgery, rats were monitored and individually handled for at least seven days before the start of behavioural experiments.

### Infusions

DL-2-Amino-5-phosphonopentanoic acid (APV, Sigma A5282) was dissolved in saline to reach a final concentration of 5 μg/ μL, pH 7.0. Clasto-Lactacystin beta-lactone (β-lac, Abcam ab141412) was first dissolved in 2% DMSO-HCl and brought to a final concentration of 32 ng/μL in saline, pH 7.4.

Antisense oligodeoxynucleotides (ODNs) were obtained from Integrated DNA Technologies at 2 nmol/μL dissolved in TE Buffer-PBS, pH 7.5, for both PKMζ-antisense and scrambled controls. The sequence for single-stranded PKMζ-antisense was C*T*C*TTGGGAAGGCAT*G*A*C and the sequence for the scrambled control was A*A*C*AATGGGTCGTCT*C*G*G where each asterisk represents a phosphorothioate linkage from 5’ to 3’. These sequences followed previous work showing selective impairment of PKMζ synthesis after PKMζ-antisense administration but not the scrambled ODN (Tsokas et al., 2016). The PKMζ-antisense ODN is the complementary sequence to start site of PKMζ mRNA whereas the scrambled control had no complementarity to a known mRNA sequence.

All infusions were performed bilaterally into the basolateral amygdala at a rate of 0.25 μL/min with a total volume of 0.5 μL/side. Intracranial infusions utilized 23-gauge injectors (Plastics One) connected to 20-gauge polyethylene tubing (Braintree Scientific, Inc.) which were connected to 26-gauge Hamilton syringes (Model 1701N). Injectors extended 1.5 mm beyond the guide cannula and remained in place for an additional one minute following the infusion to ensure proper drug diffusion. Rats were handled by the experimenter during infusions and returned to their home cage following each infusion. For experiments where infusions occurred prior to retrieval, each rat was given a sham infusion before each day of habituation to habituate rats to the infusion experience. Sham infusions followed the same procedure but no solutions were injected.

### Fear conditioning

Each day, rats were transported to a nearby holding room 30 min before the start of the experiment. Experiments utilized two different Coulbourn Habitest (model I-I10-24A) conditioning chambers referred to here as Context A and Context B. Context A had white, curved, plastic walls, and a plastic, white floor. Context B had square, checkered walls, stainless-steel grid floors, and a vanilla scent was sprayed in the chamber before each rat entered. Additionally, conditioning boxes for Context A were housed in a room of bright ambient lighting whereas Context B was in a different room with very low lighting.

In each experiment, rats were habituated and trained as follows. On Days 1 and 2, rats were placed in Context A for 20 minutes in order to habituate to the context. On Day 3, rats were placed in Context B for training. During training, rats were allowed to habituate to the context for 2 minutes followed by a 30 second tone (4 kHz, 75 dB) which co-terminated with a 1 second, 1.0 mA shock. Rats remained in the context for an additional 30 seconds before being removed. Following habituation and training:

#### Experiment 1

On Day 4, one group of rats was placed back in Context A. After 2 minutes in the box, these rats were exposed to one unpaired tone (30 seconds, 4 kHz, 75 dB) and remained in the context for an additional 30 seconds (ie. a retrieval test). Rats were returned to their home cage following this reactivation session and sacrificed 1 h later. The other group of rats did not undergo reactivation and remained in their home during this period. These non-reactivated rats were sacrificed at the same time as the reactivated group. Rats’ brains were flash frozen and collected for Western blot analysis.

#### Experiment 2

On Day 4, rats received an intracranial infusion of APV (5 μg/μL; 0.5 μL per side) or vehicle to the BLA. After 30 min, rats underwent a retrieval test (as in Experiment 1). Rats were returned to their home cage following the retrieval test and sacrificed 1 h later. Their brains were flash frozen and collected for Western blot analysis.

#### Experiment 3

On Day 4, rats underwent a retrieval test (as in Experiment 1). Immediately after retrieval, rats were infused intracranially with β-lac (32 ng/μL; 0.5 μL per side) or vehicle in the BLA. Rats were returned to their home cage following the infusion and sacrificed 1 h post-retrieval. Their brains were flash frozen and collected for Western blot analysis.

#### Experiment 4

On Day 4, rats underwent a retrieval test (as in Experiment 1). After retrieval, rats were returned to their home cage and one group was sacrificed 1 h later while the other group was sacrificed 24 h later. Rats’ brains were flash frozen and collected for Western blot analysis.

#### Experiment 5

On Day 4, rats underwent a reactivation test (as in Experiment 1). Immediately after retrieval, rats received intracranial BLA infusions with either PKMζ-antisense (2 nmol/μL; 0.5 μL per side) or scrambled control and then returned to their home cage. One day later (Day 5, PRLTM test), rats were again placed in Context A and received one unpaired tone as on Day 4. Freezing behaviour for both the retrieval/reactivation and PRLTM test days was recorded.

#### Experiment 6

Rats underwent the same procedure as in Experiment 4 except that infusions occurred 24 ours post-retrieval. As in Experiment 4, the post-infusion test occurred 24 hours after infusions.

#### Experiment 7

In this experiment, rats underwent nearly the same procedure as in Experiment 5. However, here rats received an intracranial infusion of APV (5 μg/μL; 0.5 μL per side) or vehicle 30 min prior to the beginning of the retrieval test in Context A. During each experiment, rats were recorded during training using FreezeFrame software (Actimetrics) and during tests using GeoVision GV-600 System.

### Histology

For Western blotting, rats’ brains were quickly collected and flash frozen. Rats were placed in an induction chamber containing isoflurane. Once each rat was unconscious, but still breathing, it was decapitated using a guillotine and its brain quickly retrieved. The brain was immediately submerged in a beaker containing 2-methylbutane which was seated in a container of dry ice. Once brains were frozen, they were wrapped in aluminum foil and submerged in the dry ice before final storage at −80°C.

Where rats’ brains were not required for Western blotting, rats were sacrificed in a CO_2_ induction chamber. Brains were collected and stored in 20% sucrose-formalin solution. After 48 hours, brains were sliced using a cryostat to check for proper cannula placement.

### Subcellular fractionation

Synaptosomal fractions were obtained from BLA tissue using a previously established procedure (Bai & Witzmann, 2007). Frozen brains were mounted on a cryostat and basolateral amygdala tissue was collected using a tissue puncher (Fine Science Tools). The tissue was homogenized using a Pellet Pestle (Fisher, #12141361) in 200 μL of homogenization buffer containing 20 mM HEPES, 1 mM EDTA, 2 mM EGTA, 320 mM sucrose, containing protease inhibitor (Roche, 05892791001) and phosphatase inhibitor (Roche, 04906837001) tablets. Homogenized tissue was centrifuged at 1000 g for 10 min. The supernatant was collected and centrifuged at 17,000 g for 15 min. The pellet was resuspended in 50 μL of homogenization buffer and layered on a sucrose gradient containing 100 μL of 0.8 M sucrose (1 mM EDTA, 2 mM EGTA) and 100 μL of 1.2 M sucrose (1 mM EDTA, 2 mM EGTA). This mixture was centrifuged at 54,000 g for 90 min. The layer between the 0.8 M and 1.2 M sucrose, containing the synaptosomal fraction, was collected and used for Western blotting following protein quantification with a BCA protein assay kit (Pierce).

### Western Blotting

After protein quantification, 15 μg or protein was loaded in 8% SDS-PAGE gels. Proteins were transferred overnight (at 4 °C) onto nitrocellulose membranes. Following transfer, ponceau solution (Sigma, P7170) was applied to membranes to reveal protein bands. Along with the molecular weight marker (Thermo Scientific, 26634), this enabled the cutting of membranes at 70 kDa and 40 kDa to produce two membranes. One membrane contained PKMζ protein (55 kDa) and the other contained GAPDH (37 kDa). Membranes were washed in 0.1% Tween 20, Tris-buffered saline (TBS-Tween). Blocking of membranes was done for 1 hour at room temperature in TBS-Tween containing 5% bovine serum albumin (BSA). Membranes were then incubated overnight with antibodies in 5% BSA TBS-Tween: 1:1000 PKCζ (Abcam, ab59364), 1:10,000 GAPDH (Abcam, ab8245). Following overnight incubation, membranes were washed three times with TBS-Tween. Membranes then underwent incubation with secondary antibody (Anti-rabbit IgG HRP-conjugated from Amersham, NA934V; Anti-mouse IgG HRP-conjugated from Amersham, NA931V) for 1 hour at room temperature. After secondary antibody incubation, membranes were washed four times for 10 minutes in TBS-Tween. Membranes were revealed using Pierce ECL 2 Western Blotting Substrate and scanned using a Storm Scanner (Molecular Dynamics). Scanned images were quantified using Image Lab (Bio-Rad). PKMζ protein values were compared to the GAPDH loading control for each sample. Values were then standardized as a percent of the control group.

### Statistical analyses

Data were analyzed using Jamovi (Version 1.0.7.0.). Neither homogeneity of variance, normal distribution of data, nor sphericity assumptions were violated where relevant. PKMζ protein from Western blots was analyzed using independent samples t-tests. Freezing data from behavioural experiments were analyzed using ones- or two-way repeated measures ANOVAs. Here, Tukey’s test was used for post-hoc comparisons. For all analyses, the null hypothesis was rejected where *p* < 0.05. Figures present data as means with SEM.

## Acknowledgements

We thank Josue Haubrich, Karine Gamache, Jane Zhang, and Carmelo Milo for their help with infusions. We also thank Oliver Hardt for his advice on the drafting of this manuscript and experimental design. This work was funded by grants from NSERC (RGPIN-2017-05140) and CIHR (MOP-123430). This research was undertaken thanks in part to funding from the Canada First Research Excellence Fund, awarded to McGill University for the Healthy Brains, Healthy Lives initiative.

## Author Contributions

K.N. supervised the project and designed experiments with M.B. All experiments were carried out by M.B. as well as statistical analyses and writing of the manuscript.

